## Supplementary Material for "Structural clusters of histone H3 and H4 residues regulate chronological lifespan in *Saccharomyces cerevisiae*"

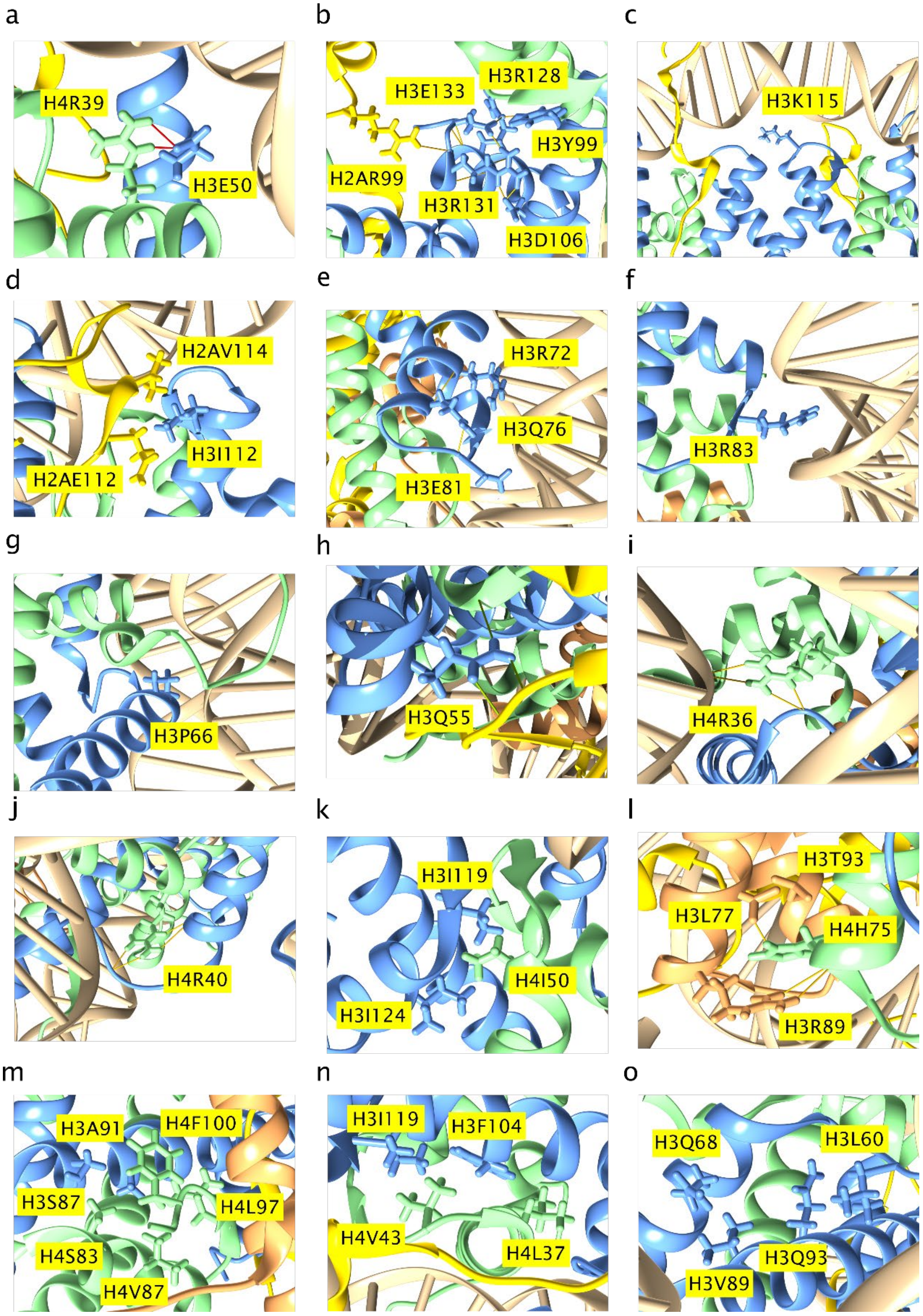

**Supplementary Fig. 1. Interactions of residues implicated in CLE.** H3E50 makes hydrogen bonds to H4R39 (**a**). H3R128 and H3R131 make H-bonds to H3Y99 and H3D106 in $\alpha$2, and H3R131 makes an H-bond to R99 in the C$\alpha$ helix of H2A (**b**). H3K115 is oriented towards a phosphate group in the DNA sugar-phosphate backbone (**c**). H3I112 forms a hydrophobic pocket with H2AQ112 and H2AV114 (**d**). H3Q76 makes bridging H-bonds to the backbone at H3D81 in L1 and H3R72 in $\alpha$1 (**e**). R83 of H3 is inserted into the minor groove at SHL ±2.5 (**f**). H3P66 at the N-terminus of $\alpha$1 may contribute to the termination the $\alpha$1 helix (**g**). H3Q55 is hydrogen bonded to the H2A C-terminal tail backbone at N110 as well as to R40 in $\alpha$1 of H4 (**h**). H4R36 make salt bridge contacts with a phosphate group in the DNA sugar-phosphate backbone (**i**). H4R40 makes H-bonds to the H3 backbone $\alpha$N and $\alpha$N-$\alpha$1 connecting coil (**j**). H4I50 is inserted between H3I119 and H3I124 (**k**). H4H75 in $\alpha$2 of H4 make hydrophobic contact with H2BL77 and H2BT93, as well as an H-bond to H2BR89 (**l**). H4F100 makes numerous hydrophobic contacts with H3S87, H3A91, H4A83, H4V87 and H4L97 (**m**). H3F104 is inserted into a hydrophobic cluster formed by H3I119, H4L37 and H4V43 (**n**). Hydrophobic contacts between H3L60 and H3Q93 and between H3Q68 and H3V89 stabilizes the conformation of H3 $\alpha$1 relative to H3 $\alpha$2 (**o**).

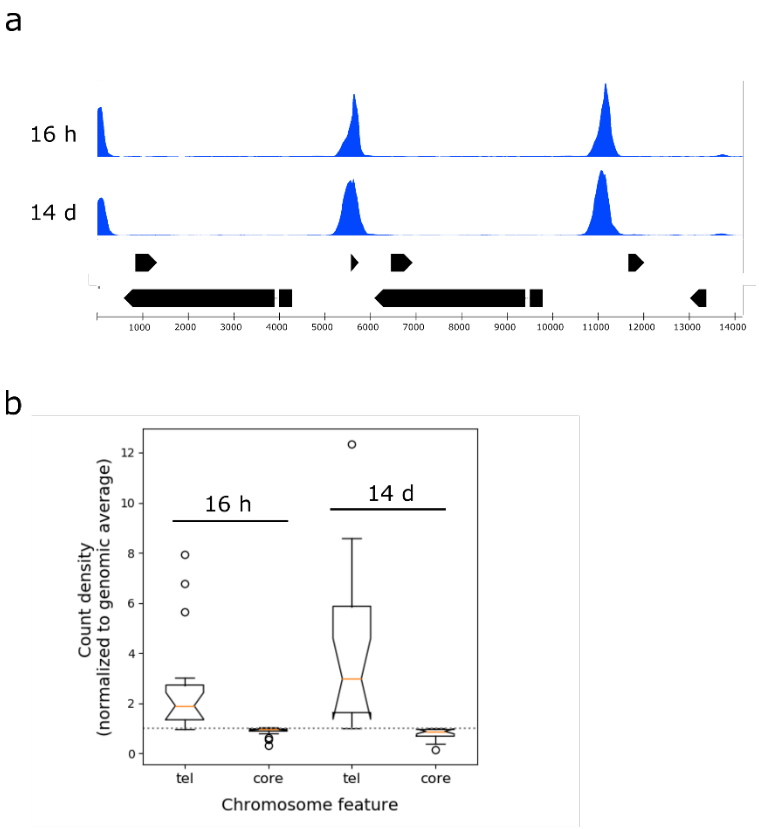


**Supplementary Fig. 2. Distribution of Rap1 in the genome during chronological aging.** The normalised binding of Rap 1 is shown at 16 h and 14 d for the left telomeric 15 kb of chromosome XII (**a**) and to the terminal 20 kb (tel) or remaining central region (core) of all 16 chromosomes normalised to the genome average (**b**). The box plots represent the inter-quartile range (IQR), the notch represents the significance at $p<0.05$, the orange line shows the median, and the whiskers is shown at 1.5× the IQR. Outliers are shown as individual data points. There is no significant difference ($p<0.05$, *t*-test) between the corresponding data groups.

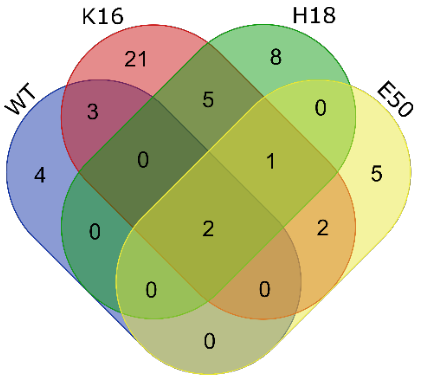


**Supplementary Fig. 3. Genes that are covered by Sir3 in different combinations of the H4K16Q, H4H18A, H3E50A and WT strains.** The Venn diagram shows the number of genes associated with Sir3 peaks that are common to various combinations of the H4K16Q, H4H18A, H3E50A and WT strains.

**Supplementary Table 1. Uploaded file "Supplementary Table 1.xlsx"**

**Supplementary Table 2. Significance of transcription factor binding sites present in redistributed Sir3 peaks.** The number of transcription factor binding sites present within assigned Sir3 peaks was determined and the statistical significance of this number $p(k)$ calculated from a Poisson distribution, $e^{-\lambda}.\frac{\lambda^k}{k!}$, where k is the number of occurrences per unit length in a Sir3 peak, and $\lambda$ is equal to the number of genomic occurrences of the site per unit length.

| Factor | Peaks | Total peaks | Total genome | Poisson pmf | Description |
|---|---|---|---|---|---|
| **H4K16Q** | | | | | |
| IME1 | 56 | 146 | 9340 | 1.94E-39 | Inducer of MEiosis |
| PDR1 | 53 | 166 | 14443 | 8.29E-29 | Pleiotropic Drug Resistance |
| STP2 | 54 | 75 | 4832 | 3.14E-21 | Protein with similarity to Stp1p |
| PUT3 | 36 | 116 | 10590 | 1.00E-19 | Proline UTilization |
| SFL1 | 33 | 66 | 36881 | 3.24E-16 | Suppressor gene for FLocculation |
| ZMS1 | 56 | 131 | 14366 | 8.96E-16 | Zinc-finger protein |
| CAD1 | 40 | 102 | 45365 | 5.40E-13 | CADmium resistance |
| NHP10 | 41 | 71 | 6479 | 9.36E-13 | Non-Histone Protein |
| TBS1 | 21 | 81 | 9241 | 1.50E-09 | ThiaBendazole Sensitive |
| OAF1 | 54 | 154 | 22341 | 3.53E-09 | Oleate-Activated transcription Factor |
| STP4 | 52 | 142 | 20357 | 8.30E-09 | Protein with similarity to Stp1p |
| NSI1 | 46 | 71 | 8375 | 3.15E-08 | NTS1 SIlencing protein 1 |
| CIN5 | 53 | 133 | 47599 | 7.24E-08 | Chromosome INstability |
| ABF1 | 67 | 141 | 21284 | 8.58E-08 | ARS-Binding Factor 1 |
| PDR3 | 16 | 34 | 3256 | 5.73E-07 | Pleiotropic Drug Resistance |
| STP3 | 47 | 113 | 17859 | 9.15E-06 | Protein with similarity to Stp1p |
| SPT15 | 24 | 73 | 25001 | 1.98E-04 | TBP basal factor |
| SWI4 | 55 | 133 | 23937 | 3.46E-04 | SWItching deficient |
| MGA1 | 37 | 67 | 22625 | 4.87E-04 | |
| STB3 | 29 | 48 | 17244 | 6.15E-04 | Sin Three Binding protein |
| PDR8 | 58 | 139 | 25911 | 7.61E-04 | Pleiotropic Drug Resistance |
| **H4H18A** | | | | | |
| IME1 | 30 | 98 | 9340 | 5.20E-22 | Inducer of MEiosis |
| PDR1 | 33 | 116 | 14443 | 1.79E-16 | Pleiotropic Drug Resistance |
| PUT3 | 21 | 87 | 10590 | 2.75E-13 | Proline UTilization |
| SFL1 | 22 | 54 | 36881 | 3.83E-13 | Suppressor gene for FLocculation |
| UGA3 | 31 | 118 | 16818 | 5.86E-13 | Utilization of GAba |

| | | | | | |
|---|---|---|---|---|---|
| REB1 | 39 | 131 | 19424 | 5.87E-13 | RNA polymerase I Enhancer Binding protein |
| CAD1 | 22 | 80 | 45365 | 1.84E-11 | CADmium resistance |
| STP2 | 32 | 47 | 4832 | 1.71E-10 | Protein with similarity to Stp1p |
| OAF1 | 37 | 131 | 22341 | 2.83E-09 | Oleate-Activated transcription Factor |
| NHP10 | 26 | 54 | 6479 | 3.74E-09 | Non-Histone Protein |
| ZMS1 | 33 | 91 | 14366 | 1.04E-08 | ReSpiration Factor |
| RPH1 | 39 | 137 | 25055 | 5.19E-08 | Regulator of PHR1 |
| STP4 | 34 | 116 | 20357 | 6.45E-08 | Protein with similarity to Stp1p |
| MET31 | 37 | 145 | 27447 | 1.36E-07 | METhionine requiring |
| NSI1 | 31 | 58 | 8375 | 2.53E-07 | NTS1 SIlencing protein 1 |
| GAL4 | 37 | 122 | 22321 | 2.87E-07 | GALactose metabolism |
| CIN5 | 38 | 110 | 47599 | 3.09E-06 | Chromosome INstability |
| SPT15 | 16 | 48 | 25001 | 4.01E-06 | SuPpressor of Ty insertions |
| TBS1 | 14 | 56 | 9241 | 1.59E-05 | ThiaBendazole Sensitive |
| STP3 | 30 | 93 | 17859 | 1.77E-05 | Protein with similarity to Stp1p |
| RAP1 | 36 | 120 | 24608 | 2.62E-05 | Repressor/Activator site binding Protein |
| ABF1 | 38 | 96 | 21284 | 0.00109 | ARS-Binding Factor 1 |

**Supplementary Table 3. Annotated genes that are associated with elevated genomic Sir3 levels in WT, H4K16Q, H4H18A and H3E50A yeast strains.**

| Strain(s) in group | Number genes shared | Genes shared |
|---|---|---|
| **E50 H18 K16 WT** | 2 | YFR034C YFR033C |
| **E50 H18 K16** | 1 | YIL055C |
| **K16 WT** | 3 | YFR018C YEL035C YFR017C |
| **H18 K16** | 5 | YOL060C YLR227W-A YDR186C YLR227W-B YEL008C-A |
| **E50 K16** | 2 | YMR194C-B YOR348C |
| **WT** | 4 | YMR142C YCL069W YCL068C YCR096C |
| **K16** | 21 | YOR375C YGR038C-B YDR506C YEL060C YMR069W YNL104C YIL047C YEL036C YLR141W YGL126W YJL027C YFL016C YFL015C YDR488C YML101C YDR505C YJR113C YGR038C-A YDR487C YOR301W YKL097C |
| **H18** | 8 | YGR027W-B YGR027W-A YOL110W YKL029C YOL111C YCL025C YGR050C YOL012C |
| **E50** | 5 | YCR006C YPL171C YJL133C-A YKR096W YCR005C |

**Supplementary Table 4. GO biological function categories that show significant enrichment for genes that are induced (adjusted p<0.05) in the H4K16Q, H4H18A and H3E50A mutant strains compared to the WT strain.** The reported probability value is calculated as the probability mass function of a hypergeometric distribution using the values in the Enrichment column, and the q-value is the Benjamini–Hochberg corrected p-value.

| GO biological function | Probability (p) | FDR q-value | Enrichment (N, K, n, k) |
|---|---|---|---|
| **H4K16Q mutant** | | | |
| **Cytoplasmic translation** | 5.68E-59 | 2.99E-55 | 3.75 (5768,132,1376,118) |
| **Organonitrogen compound biosynthetic process** | 1.08E-54 | 2.85E-51 | 2.02 (5768,736,1376,355) |
| **Ribonucleoprotein complex biogenesis** | 1.57E-49 | 2.75E-46 | 2.93 (5768,222,1376,155) |
| **Ribosome biogenesis** | 3.76E-45 | 4.94E-42 | 2.97 (5768,195,1376,138) |
| **Cellular component biogenesis** | 1.91E-44 | 2.01E-41 | 2.72 (5768,247,1376,160) |
| **ncRNA metabolic process** | 2.40E-44 | 2.10E-41 | 2.17 (5768,482,1376,250) |
| **rRNA processing** | 6.01E-44 | 4.52E-41 | 2.66 (5768,258,1376,164) |
| **ncRNA processing** | 8.06E-42 | 5.30E-39 | 2.31 (5768,375,1376,207) |
| **Amide biosynthetic process** | 1.55E-38 | 9.05E-36 | 2.27 (5768,370,1376,200) |
| **rRNA metabolic process** | 6.73E-38 | 3.54E-35 | 2.41 (5768,301,1376,173) |
| **H4H18A mutant** | | | |
| **Organonitrogen compound biosynthetic process** | 7.38E-79 | 3.88E-75 | 2.69 (5768,736,899,309) |
| **Cytoplasmic translation** | 4.60E-78 | 1.21E-74 | 5.64 (5768,132,899,116) |
| **Peptide biosynthetic process** | 1.78E-58 | 3.12E-55 | 3.40 (5768,321,899,170) |
| **Translation** | 2.08E-58 | 2.73E-55 | 3.41 (5768,318,899,169) |
| **Amide biosynthetic process** | 4.00E-57 | 4.21E-54 | 3.17 (5768,370,899,183) |
| **Peptide metabolic process** | 1.67E-52 | 1.46E-49 | 3.14 (5768,352,899,172) |
| **Cellular biosynthetic process** | 1.61E-50 | 1.21E-47 | 1.88 (5768,1346,899,394) |
| **Cellular amide metabolic process** | 3.24E-50 | 2.13E-47 | 2.83 (5768,435,899,192) |
| **Organic substance biosynthetic process** | 3.55E-50 | 2.08E-47 | 1.86 (5768,1386,899,401) |
| **Biosynthetic process** | 3.78E-50 | 1.99E-47 | 1.85 (5768,1402,899,404) |
| **H3E50A mutant** | | | |
| **Oxidation-reduction process** | 2.30E-19 | 1.21E-15 | 1.93 (5768,410,1136,156) |
| **Generation of precursor metabolites and energy** | 2.07E-16 | 5.46E-13 | 2.46 (5768,157,1136,76) |
| **Antibiotic metabolic process** | 1.06E-15 | 1.86E-12 | 3.33 (5768,64,1136,42) |
| **Energy derivation by oxidation of organic compounds** | 1.70E-15 | 2.23E-12 | 2.82 (5768,99,1136,55) |
| **Tricarboxylic acid cycle** | 9.40E-14 | 9.89E-11 | 4.23 (5768,30,1136,25) |
| **Citrate metabolic process** | 9.40E-14 | 8.24E-11 | 4.23 (5768,30,1136,25) |
| **Electron transport chain** | 3.17E-13 | 2.38E-10 | 3.26 (5768,56,1136,36) |

| | | | |
|---|---|---|---|
| **Respiratory electron transport chain** | 3.61E-13 | 2.37E-10 | 4.00 (5768,33,1136,26) |
| **Tricarboxylic acid metabolic process** | 3.95E-13 | 2.31E-10 | 4.09 (5768,31,1136,25) |
| **Cellular respiration** | 3.13E-11 | 1.65E-08 | 3.03 (5768,57,1136,34) |

**Supplementary Table 5. KEGG pathways associated with genes that are significantly upregulated (adjusted $p<0.05$) in the H4K16Q, H4H18A and H3E50A mutant compared to the WT strain.** *S. cerevisiae* KEGG pathways were identified with KEGG Mapper[1], and are arranged by descending mapping frequency.

| Pathway | Description | Number of genes |
|---|---|---|
| | **H4K16Q** | |
| **sce01100** | Metabolic pathways | 263 |
| **sce01110** | Biosynthesis of secondary metabolites | 137 |
| **sce03010** | Ribosome | 136 |
| **sce01130** | Biosynthesis of antibiotics | 118 |
| **sce01230** | Biosynthesis of amino acids | 86 |
| **sce01200** | Carbon metabolism | 50 |
| **sce03008** | Ribosome biogenesis in eukaryotes | 47 |
| **sce03013** | RNA transport | 44 |
| **sce00230** | Purine metabolism | 29 |
| **sce01210** | 2-Oxocarboxylic acid metabolism | 26 |
| | **H4H18A** | |
| **sce01100** | Metabolic pathways | 202 |
| **sce03010** | Ribosome | 131 |
| **sce01110** | Biosynthesis of secondary metabolites | 115 |
| **sce01130** | Biosynthesis of antibiotics | 98 |
| **sce01230** | Biosynthesis of amino acids | 69 |
| **sce01200** | Carbon metabolism | 35 |
| **sce00230** | Purine metabolism | 28 |
| **sce03013** | RNA transport | 27 |
| **sce00010** | Glycolysis / Gluconeogenesis | 23 |
| **sce03008** | Ribosome biogenesis in eukaryotes | 22 |
| | **H3E50A** | |
| **sce01100** | Metabolic pathways | 195 |
| **sce01110** | Biosynthesis of secondary metabolites | 78 |
| **sce01130** | Biosynthesis of antibiotics | 60 |
| **sce01200** | Carbon metabolism | 49 |
| **sce00190** | Oxidative phosphorylation | 38 |
| **sce00620** | Pyruvate metabolism | 27 |
| **sce00020** | Citrate cycle (TCA cycle) | 26 |
| **sce04138** | Autophagy | 24 |
| **sce01230** | Biosynthesis of amino acids | 20 |
| **sce04146** | Peroxisome | 20 |

[1]Kanehisa, M., Goto, S., Sato, Y., Furumichi, M. & Tanabe, M. KEGG for integration and interpretation of large-scale molecular data sets. Nucleic Acids Res. 40, D109-14 (2012).

**Supplementary Table 6. GO biological function categories that show significant enrichment for genes that are repressed (adjusted p<0.05) in the H4K16Q, H4H18A and H3E50A strains compared to the WT strain.** The reported probability value is calculated as the probability mass function of a hypergeometric distribution using the values in the Enrichment column, and the q-value is the Benjamini–Hochberg corrected p-value.

| GO biological function | Probability (p) | FDR q-value | Enrichment (N, K, n, k) |
|---|---|---|---|
| **H4K16Q** | | | |
| **DNA integration** | 1.41E-14 | 7.40E-11 | 4.02 (5768,49,938,32) |
| **DNA recombination** | 1.55E-14 | 4.08E-11 | 2.31 (5768,213,938,80) |
| **Reproductive process** | 5.97E-11 | 1.05E-07 | 1.76 (5768,401,938,115) |
| **Meiotic cell cycle** | 7.45E-11 | 9.80E-08 | 2.41 (5768,135,938,53) |
| **RNA-dependent DNA biosynthetic process** | 9.10E-11 | 9.57E-08 | 3.07 (5768,68,938,34) |
| **Meiotic cell cycle process** | 6.52E-09 | 5.71E-06 | 1.89 (5768,244,938,75) |
| **Transposition** | 2.56E-07 | 1.93E-04 | 2.30 (5768,99,938,37) |
| **DNA biosynthetic process** | 3.38E-07 | 2.22E-04 | 2.31 (5768,96,938,36) |
| **Transposition, RNA-mediated** | 4.45E-07 | 2.60E-04 | 2.31 (5768,93,938,35) |
| **Mitochondrial respiratory chain complex assembly** | 2.55E-06 | 1.34E-03 | 2.81 (5768,46,938,21) |
| **H4H18A** | | | |
| **DNA integration** | 8.96E-13 | 4.71E-09 | 3.08 (5768,49,1338,35) |
| **DNA recombination** | 1.78E-12 | 4.68E-09 | 1.92 (5768,213,1338,95) |
| **RNA-dependent DNA biosynthetic process** | 8.93E-12 | 1.57E-08 | 2.66 (5768,68,1338,42) |
| **DNA biosynthetic process** | 2.04E-09 | 2.68E-06 | 2.20 (5768,96,1338,49) |
| **Cell cycle process** | 1.23E-07 | 1.29E-04 | 1.39 (5768,571,1338,184) |
| **DNA metabolic process** | 1.39E-07 | 1.22E-04 | 1.43 (5768,480,1338,159) |
| **Meiotic cell cycle** | 1.36E-06 | 1.02E-03 | 1.79 (5768,135,1338,56) |
| **Cell cycle** | 1.21E-05 | 7.95E-03 | 1.38 (5768,408,1338,131) |
| **Meiotic cell cycle process** | 1.69E-05 | 9.90E-03 | 1.50 (5768,244,1338,85) |
| **Reproductive process** | 2.10E-05 | 1.11E-02 | 1.38 (5768,401,1338,128) |
| **H3E50A** | | | |
| **Ribonucleoprotein complex biogenesis** | 1.67E-47 | 8.81E-44 | 2.93 (5768,222,1341,151) |
| **Cellular component biogenesis** | 6.97E-47 | 1.83E-43 | 2.80 (5768,247,1341,161) |
| **Ribosome biogenesis** | 6.73E-43 | 1.18E-39 | 2.96 (5768,195,1341,134) |
| **Cytoplasmic translation** | 3.21E-40 | 4.22E-37 | 3.32 (5768,132,1341,102) |
| **rRNA processing** | 2.00E-36 | 2.10E-33 | 2.53 (5768,258,1341,152) |
| **rRNA metabolic process** | 1.68E-33 | 1.48E-30 | 2.34 (5768,301,1341,164) |
| **ncRNA metabolic process** | 6.41E-29 | 4.81E-26 | 1.95 (5768,482,1341,218) |
| **ncRNA processing** | 1.11E-28 | 7.30E-26 | 2.09 (5768,375,1341,182) |

| | | | |
|---|---|---|---|
| **Maturation of SSU-rRNA** | 8.15E-22 | 4.76E-19 | 3.17 (5768,80,1341,59) |
| **maturation of SSU-rRNA from tricistronic rRNA transcript (SSU-rRNA, 5.8S rRNA, LSU-rRNA)** | 4.24E-21 | 2.23E-18 | 3.34 (5768,67,1341,52) |

**Supplementary Table 7. KEGG pathways associated with genes that are repressed (adjusted p<0.05) in the H4K16Q, H4H18A and H3E50A mutant compared to the WT strain**. *S. cerevisiae* KEGG pathways were identified with KEGG Mapper, and are arranged by descending mapping frequency.

| Pathway | Description | Number of genes (of 844) |
|---|---|---|
| | **H4K16Q** | |
| **sce01100** | Metabolic pathways | 73 |
| **sce04113** | Meiosis | 37 |
| **sce01110** | Biosynthesis of secondary metabolites | 29 |
| **sce04111** | Cell cycle | 27 |
| **sce04011** | MAPK signaling pathway | 22 |
| **sce01130** | Biosynthesis of antibiotics | 20 |
| **sce04138** | Autophagy | 14 |
| **sce01200** | Carbon metabolism | 13 |
| **sce03010** | Ribosome | 11 |
| **sce04144** | Endocytosis | 10 |
| | **H4H18A** | |
| **sce01100** | Metabolic pathways | 108 |
| **sce04113** | Meiosis | 39 |
| **sce04111** | Cell cycle | 36 |
| **sce01110** | Biosynthesis of secondary metabolites | 33 |
| **sce04011** | MAPK signaling pathway | 28 |
| **sce01130** | Biosynthesis of antibiotics | 25 |
| **sce04138** | Autophagy | 23 |
| **sce03040** | Spliceosome | 19 |
| **sce01200** | Carbon metabolism | 16 |
| **sce04146** | Peroxisome | 15 |
| | **H3E50A** | |
| **sce01100** | Metabolic pathways | 192 |
| **sce03010** | Ribosome | 123 |
| **sce01110** | Biosynthesis of secondary metabolites | 77 |
| **sce01130** | Biosynthesis of antibiotics | 65 |
| **sce01230** | Biosynthesis of amino acids | 41 |
| **sce04111** | Cell cycle | 40 |
| **sce03008** | Ribosome biogenesis in eukaryotes | 39 |
| **sce04113** | Meiosis | 33 |
| **sce04011** | MAPK signaling pathway | 33 |
| **sce00230** | Purine metabolism | 27 |

**Supplementary Note**

### Analysis of structure and interactions of residues implicated in chronological lifespan

#### Residues implicated in extension of lifespan

To understand the impact of mutating the residues at the histone fold fringes on nucleosome structure, we identified the interactions of the wild type residues at each position in the nucleosome. In the H3 C-terminal region of $\alpha$3, R128 and R131 make H-bonds to H3Y99 and H3D106 in the central region of $\alpha$2, and H3R131 makes an H-bond to R99 in the C$\alpha$ helix of H2A (Supplementary Fig. 1a). These interactions probably stabilise the position of the C-terminus of H3 in the nucleosome and contribute to the formation of a stable four helix bundle connecting the two H3-H4 dimers. H3K115 is oriented towards a phosphate group in the DNA sugar-phosphate backbone, and H3I112 forms a hydrophobic cluster with H2AQ112 and H2AV114 (Supplementary Fig. 1b,c). H3Q76 makes bridging H-bonds to the backbone at H3D81 in L1 and H3R72 in $\alpha$1, which may stabilize the L1 loop conformation (Supplementary Fig. 1d). Whereas substitution of H3Q76 with E extends lifespan, substitution with A reduces lifespan to less than the median of the population (Supplementary Table 1). H3R83 is inserted into the minor groove at position SHL ±2.5 (Supplementary Fig. 1e). Arginine is inserted into the minor groove at six other positions at SHL ±6.5 (H3R49), ±5.5 (H2AR47) and ±0.5 (H4R45), giving a total of 8 such insertions over the length of the nucleosome DNA duplex[1]. H3P66 at the N-terminus of $\alpha$1 may contribute to the termination the $\alpha$1 helix and formation of a coil between the $\alpha$N and $\alpha$1 helices of H3 (Supplementary Fig. 1f). In the region of the structure where H4 $\alpha$1 passes over H3 $\alpha$N, H3Q55 is hydrogen bonded to the H2A C-terminal tail backbone at N110 as well as to R40 in H4 $\alpha$1 (Supplementary Fig. 1g). H4R36 is oriented towards the DNA, and makes a salt bridge with a backbone phosphate (Supplementary Fig. 1h). As expected, the charge of the residue is crucial, since whereas the R→K substitution extends lifespan, the R→A change reduced lifespan to less than the population median (Supplementary Table 1). H4R40 makes H-bonds to the backbone at positions in the C-terminus of $\alpha$N and the coil connecting $\alpha$N and $\alpha$1 of H3 (Supplementary Fig. 1i). I50 at the N-terminus of H4 $\alpha$2 is inserted between H3I119 and H3I124, allowing hydrophobic contact, and fixing the position of the N-terminus of H4 $\alpha$2 to $\alpha$3 of H3 (Supplementary Fig. 1j). H75 in the C-terminus of H4 $\alpha$2 make hydrophobic contact with L77 in the $\alpha$2 C-terminus and T93 in the centre of $\alpha$3 of H2B, as well as H-bonds to R89 at the N-terminus of H2B $\alpha$3 (Supplementary Fig. 1k). These interactions will contribute to the stability of the four helix bundle that binds a H3-H4 dimer to a H2A-H2B dimer. F100 in the short C-terminal tail of H4 makes numerous hydrophobic contacts with A83 and V87 in the $\alpha$3 of H4, as well as L97 in the C-terminal $\beta$-strand. Additional contacts are also seen between H4F100 and S87 and A91 in the N-terminal part of the H3 $\alpha$2 (Supplementary Fig. 1l).

All these interactions, apart from being concentrated at the extremities of the histone fold domains of H3 and H4, involve interactions either with other dimers, interactions between the αN and α1 domains of H3 and H4, or interactions with the DNA duplex. The interactions specifically exclude the extensive interactions between dimer partners, and seem enriched for contacts that stabilize the H3-H4 dimer in its frame within the nucleosome structure.

**Residues implicated in reduction of lifespan**

H3F104 interleaves with a cluster of hydrophobic residues composed of H4L37, H4V43 and H3I119 (Supplementary Fig. 1n), and is likely to contribute to the stable placement of the C-terminal region of the α2 helix and the L2 loop of H3 relative to the α1 helix and L1 loop of H4. H3L60 in the coiled region, joining the Nα and α1 helices, approaches H3Q93 in the α2 loop, and H3Q68 in α1 approaches α2 H3V89, setting the path of the H3 tail to a trajectory between the DNA gyres (Supplementary Fig. 1o). H4D85 is hydrogen bonded to H4R78 and H4T72, possibly stabilizing the tight turn of L2 between α2 and α3. We speculate that the mutations that cause a shortened lifespan contribute to the structural destabilization of the histone octamer and disruption of chromatin structures.
